# Accelerate the discovery of genetic variants in mitochondrial diseases with VIOLA: Variant PrIOritization using Latent space

**DOI:** 10.1101/2025.05.16.654430

**Authors:** Justine Labory, Youssef Boulaimen, Jasmine Singh, Samira Ait-el- Mkadem Saadi, Véronique Paquis-Flucklinger, Sylvie Bannwarth, Silvia Bottini

## Abstract

Interpreting variants from whole-exome sequencing remains a major challenge, particularly for heterogeneous disorders such as mitochondrial diseases (MD). To help finding a diagnosis for complex cases, we have developed VIOLA (Variant prIoritizatiOn using Latent spAce). The pipeline includes a variational autoencoder to embed functional annotations into a low-dimensional space, followed by DBSCAN-based outlier detection. To prioritize potential pathogenic variants, filtering steps and phenotype integration via HPO terms are then applied. Two scores are associated to the selected variants: the VIOLA score (Vscore), which combines variant features, transcriptomic co-expression data, and MD-specific annotations, and the VIOLA Aggregated score (VAscore) that merges Vscore with Exomiser’s pathogenicity score. Finally, we also provide the ARrank, specifically designed for variants compatible with autosomal recessive inheritance. We illustrate the functionalities of VIOLA on a novel cohort of patients with suspected MD with complex phenotypes. VIOLA systematically ranked causal variants among the top, outperforming existing methods. Overall, VIOLA is a patient-specific approach to help to discover novel variants in complex MD.

## Introduction

Rare diseases are defined as conditions affecting fewer than one in 2,000 individuals, but collectively they impact millions of people worldwide^1^. Despite their individual rarity, there are over 7,000 rare diseases^2^, and the majority have a genetic cause. Diagnosis of these diseases is often a complex process, often referred to as a “diagnostic odyssey”, which can take years and involves numerous clinical assessments and tests. One of the main challenges in diagnosing rare diseases is identifying the pathogenic genetic variants responsible for these conditions’ basis^3^.

Among the most difficult to diagnose, there are the mitochondrial diseases (MD). Those are genetic rare disorders caused by the deficiency of the mitochondrial respiratory chain, which provides energy in each cell through oxidative phosphorylation^4^. The extreme heterogeneity on both clinical and genetical sides, with a broad range of age onset and very different symptoms, contribute to the difficulty in the diagnosis^5^. Mitochondrial dysfunctions can be classified into two broad categories: primary mitochondrial diseases (PMDs) and secondary mitochondrial dysfunctions (SMDs), a distinction that is clinically important but often difficult to establish^6^. PMDs are caused by germline mutations in either mitochondrial DNA (mtDNA) or nuclear DNA (nDNA) that directly impair oxidative phosphorylation^7^. SMDs, on the other hand, result from mutations in genes that do not directly code for mitochondrial proteins but whose dysfunction leads to impaired mitochondrial activity^8^. The increasing recognition of SMDs adds complexity to the diagnostic process, as they often present with symptoms overlapping those of PMDs and may show similar biochemical and clinical profiles^7^.

The modes of transmission represent another level of complexity. The mtDNA is exclusively maternally transmitted^9^, while the nuclear genes follow Mendelian inheritance, meaning that they can be transmitted in an autosomal dominant, autosomal recessive (AR) or even X-linked mode, which further complicates diagnosis and variant interpretation^10^. So, all modes of genetic transmission may be involved but 80% of the genes already known to be involved in MD follow an AR inheritance^11^.

The advent of high-throughput genomic sequencing technologies, such as whole genome sequencing (WGS) and whole exome sequencing (WES), has dramatically increased our ability to detect genetic variants^12, 13^. However, the big number of variants identified in a single genome, typically in the tens of thousands, poses a major challenge when it comes to distinguishing those that are pathogenic from the huge majority that are benign. This identification process is crucial, as finding the causal variant can lead to accurate diagnosis, therapeutic decisions and better disease management. However, identifying the causal variant is time-consuming and require experts’ curation^14^. This is why the use of variant prioritization becomes essential. Variant prioritization is the process of assigning priority or rank to genetic variants identified in a genomic analysis based on their potential clinical significance or relevance. This method aims to identify variants that are most likely to be pathogenic or have a significant phenotypic impact on an individual. Conventional approaches to variant prioritization rely on amino acid or nucleotide conservation to predict the pathogenicity of missense variant and the impacts on protein function^15, 16^. These approaches are limited by predefined rules and existing knowledge. Recent variant prioritization tools leverage machine and deep learning techniques^17–20^. These techniques can analyze vast amounts of genomics data, identify complex patterns, and make predictions based on these patterns. Among those, an outstanding example is CADD (Combined Annotation Dependent Depletion), which integrates multiple annotations into a single score to estimate the deleteriousness of variants across the genome by using support vector machine (SVM)^17, 21^ and Exomiser uses a logistic regression model to combine variant-based and gene-based scores to generate a final combined score used for ranking^18^. Although Exomiser is nowadays considered as “state-of-the art” variant prioritization tool, it is not tailored for MD cases where responsible variant(s) usually is characterized by a peculiar combination of parameters not found in other patients/diseases. A particularly promising area within machine learning for uncovering non-linear relationships is the use of variational autoencoders (VAEs). VAEs are generative models that perform dimensionality reduction by learning latent representations of high-dimensional data. VAEs model the underlying data distribution probabilistically, making them well-suited for capturing non-linear relationships.

Here, we use a VAE to identify non-linear relationship among variant features to improve variant prioritization in the context of MD. Therefore, we introduce VIOLA (Variant PrIOritization using Latent spAce), a novel approach based on the innovative hypothesis that since the disease-responsible variant(s) are individual-specific and rare, therefore they should have a unique combination of properties different from the rest of the individual variants. Hence, we assume that the putative disease variants for MD suspicion are outliers of each individual variants’ distribution. Based on this hypothesis, VIOLA first collects 32 scores spanning from pathogenicity to splicing to describe variants properties. Then uses a VAE to transform those properties in a set of latent features. At the level of the latent space, a density-based clustering technique is used to retrieve the outliers’ variants. The remaining variants are filtered according to quality criteria and patient phenotype. Finally, VIOLA provides two scores: the VIOLA score (Vscore) obtained by combining Mahalanobis distance metric, transcriptomics data and MD-related features and the VIOLA Aggregated score (VAscore) which integrates Exomiser score.

## Materiel and Methods

### Patient cohort

We dispose of WES and RNA-sequencing data from 20 patients with suspicion of MD, of which 4 with known responsible variant. The demographic details of the cohort are presented in Table I. For 4 patients, 3 causative genes were identified (the same gene for 2 patients since they are cousins), two of them were reported in Clinvar as pathogenic. The main characteristics of these variants can be found in Table II.

**Table I:** Characteristics of the cohort. Description of the characteristics of the patient cohort, including anonymized patient IDs, sex, and age at sample collection.

| Patient id | Sex | Age at disease onset | Age at skin biopsy |
| --- | --- | --- | --- |
| 1 | Male | 2 weeks | 2 weeks |
| 2 | Male | 24 years old | 26 years old |
| 3 | Female | 24 hours | 24 hours |
| 4 | Female | < 1 year old | 2 years old |
| 5 | Female | 18 years old | 27 years old |
| 6 | Male | Adult | 70 years old |
| 7 | Male | Adult | 60 years old |
| 8 | Male | < 1 year old | 2 months |
| 9 | Female | 2 years old | 4 years old |
| 10 | Male | 5 years old | 11 years old |
| 11 | Female | 5 years old | 5 years old |
| 12 | Male | 3 months | 8 months |
| 13 | Male | 17 years old | 19 years old |
| 14 | Male | 50 years old | 56 years old |
| 15 | Male | 9 months | 6 years old |
| 16 | Male | 9 years old | 9 years old |
| 17 | Male | 4 months | 19 months |
| 18 | Female | < 1 year old | 11 months old |
| 19 | Male | 18 months | 15 years old |
| 20 | Female | 2 months | 8 years old |

**Table II:** Characteristics of the responsible variants. Overview of the characteristics of the responsible variants for mitochondrial diseases, including their type, chromosomal location, affected gene, and predicted functional consequence by Variant Effect Predictor for the four diagnosed patients.

| Patient | Variant type | Chromosome | Position | Gene | Consequence |
| --- | --- | --- | --- | --- | --- |
| 3 | Deletion | 13 | 38349764 | UFM1 | Upstream |
| 4 | SNV | 1 | 12308619 | VPS13D | Stop gained |
| 9 | SNV | 3 | 123352464 | ADCY5 | Missense |
| 17 | Deletion | 13 | 38349764 | UFM1 | Upstream |

In patients 3 and 17, we found the NM_016617.4:c.-155_-153del (or NM_001286704.2: c.-273_-271del) in the *UFM1* gene which was homozygous in the two patients. This variant is in the promoter region and was reported in ClinVar as a founder mutation causing early-onset severe encephalopathy^22^ (ClinVar variation ID: 495149), similar to that of our patients’ symptoms. They presented a severe encephalopathy with failure to thrive, dystonia, severe developmental delay, and myelination abnormality. The *UFM1* gene encodes ubiquitin-fold modifier 1 (UFM1), a ubiquitin-like (UBL) protein, which is ubiquitously expressed, including in brain.

In patient 9, the heterozygous *de novo* variant NM_183357.2:c.1252C>T, p.Arg418Trp in the *ADCY5* gene, reported as pathogenic and responsible of an autosomal dominant Dyskinesia with orofacial involvement^23^ (ClinVar variation ID: 162090), was identified in our patient with connected symptoms: axial hypotonia, dyskinesia and dystonia and increased lactate and mitochondrial proliferation.

Finally, for patient 4, the responsible variant identified was the NM_015378.3:c.6628C>T p.Gln2210Ter in the *VPS13D* gene. It was a loss of function variant (nonsense), very rare in population databases, and reported as ‘likely pathogenic’ in ClinVar (ClinVar variation ID: 3767166)^24^. *VPS13D* biallelic pathogenic variants have been reported in patients displaying variable neurological phenotypes, with an autosomic recessive inheritance. *VPS13D* movement disorder is a hyperkinetic movement disorder (dystonia, chorea, and/or ataxia) of variable age of onset that can be associated with developmental delay. Onset ranges from birth to adulthood. Clinical presentation of infantile-onset is characterized by hypotonia and severe developmental delay or motor delay that progresses to severe generalized dystonia or spastic ataxia. The phenotype was comparable to that of our patient. After an intrauterine growth retardation, she presented in neonatal period with vomiting, microcephaly, seizure, increased lactate, basal ganglia abnormalities^24^.

Unlike *VPS13D*, a known mitochondrial gene already associated with mitochondrial diseases, *UFM1* and *ACY5* are not classified as mitochondrial genes, and their connection to mitochondrial function has yet to be established.

#### Genomics data

DNA was extracted from whole blood and exonic regions were enriched using the SureSelect Human All Exon kit from Agilent followed by sequencing as 75□bp paired-end runs on an Illumina HiSeq2000 (samples 1 to 5), 100 paired-end runs on Illumina HiSeq2500 (samples 6 to 9) or 75□bp paired-end runs on HiSeq4000 (samples 10 to 20). Raw image files were processed by the IlluminaReal Time Analysis pipeline for base calling and generating the read sets.

#### Transcriptomics data

Non-strand specific, polyA-enriched RNA-seq was performed as follows: RNA was isolated from patient fibroblasts lysates using the AllPrep RNA Kit (Qiagen) and RNA integrity number (RIN) were determined with the Agilent 2100 BioAnalyzer (RIN>7). RNA quantification and quality control were performed by PCR and BioAnalyzer profiling. For library preparation, 1 µg of RNA was poly(A) selected, fragmented and reverse transcribed with the Elute, Prime and Fragment Mix (Illumina). End repair, A-tailing, adaptor ligation and library enrichment was performed as described in the protocol of the ‘TruSeq Stranded mRNA Sample Prep’ of Illumina. Libraries were assessed for quality and quantity with the Agilent 2100 BioAnalyzer and PCR. Libraries were sequenced as 2x75 nucleotides paired end runs on an Illumina HiSeq4000 platform. For each sample, 50 million of clusters and 100 million of sequences (7.5Gb) +/-12% were generated (Integragen, France).

### Bioinformatic analysis of genomics data to identify individuals’ variants

#### Step1: Variant Calling

Raw sequence reads underwent quality control (QC) with FastQC tool^25^ and were mapped to the GRCh38 reference genome with GATK^26^. Variants were subsequently called in the form of single nucleotide variants (SNVs) and small insertions/deletions (indels) with HaplotypeCaller. The resulting variant calling format (VCF) files were filtered using hard filter settings. We kept default parameters except for the filter concerning StrandOddsRatio that we removed because for some patients of the cohort, the responsible variant was filtered out because of this filter.

#### Step 2: Variant annotation

We used the tool Variant Effect Predictor (VEP) to annotate genetic variants^27^, providing their genomics locations and their potential effects on genes, transcripts, and proteins. The tool annotated each variant with key information, including gene names, transcript IDs, variant consequence predictions (such as missense, nonsense, or synonymous changes), and known pathogenicity from databases like ClinVar^28, 29^ or dbSNP^30^. VEP also provided allele frequencies from Genome Aggregation Database (gnomAD)^31^ and 1000 Genomes (1000G)^32^ allowing us to filter variants based on their minor allele frequency (MAF).

#### Step 3: Variant frequency filter

We applied stringent filtering criteria on all patient variants, retaining only those variants that have a MAF less than 0.01 across gnomAD^31^ and 1000G databases^32^ therefore assuring to retrieve only rare variants. This step minimizes the inclusion of variants that might be benign or unrelated to the disease. Importantly, we also preserved variants for which frequency information are unavailable. Variants without recorded frequencies could represent extremely rare or novel mutations that have not yet been captured in existing databases. These unrecorded variants may be of particular significance in the context of rare diseases, as they could hold clues to pathogenic mechanisms or provide insights into undiagnosed cases.

### Bioinformatic analysis of transcriptomics data to identify individual variants

Quality checks were first performed on the raw RNA-seq reads using FastQC^25^ to assess sequencing quality. Then, the reads were aligned to the human genome (hg38 assembly, release 26) using the tool STAR (Spliced Transcripts Alignment to a Reference)^33^ in two-pass mode. This modality guarantees spliced alignment accuracy by performing an initial mapping step to detect novel splice junctions, which are then incorporated into the second step for a more accurate alignment.

Following alignment, duplicate reads were identified and removed using the MarkDuplicates tool from the Picard suite to minimize biases introduced by PCR amplification. The resulting BAM files were then used as input for the RNA-SeQC tool^34^, which generated read count data for each gene, ultimately producing a gene expression matrix for downstream analyses.

### MitoBook: a catalog of co-expressed genes modules in MD context

#### Co-expression analysis

The raw read counts obtained from RNA-seq analysis were used as the input for a co-expression analysis aimed at uncovering patterns of gene expression. To achieve this, we employed Weighted Gene Co-expression Network Analysis (WGCNA)^35^, a widely used method for constructing gene co-expression networks. WGCNA identifies groups of highly correlated genes, referred to as “modules,” which share similar expression profiles across the samples. These modules often represent functionally related groups of genes or biological pathways, making them valuable for understanding the molecular mechanisms underlying complex conditions such as mitochondrial diseases.

#### Enrichment in mitochondrial genes

Once the gene modules were defined, we investigated their relevance to mitochondrial biology by assessing their enrichment in mitochondrial genes using the MitoCarta database^36^. MitoCarta is a comprehensive resource that catalogues 1136 human genes encoding proteins with known or predicted mitochondrial localization. To determine whether a module was enriched in mitochondrial genes, we calculated a “propensity score” representing the proportion of MitoCarta genes within each module. To establish a threshold for enrichment, we used the module with the highest number of genes, as such modules are typically non-specific and serve as a baseline for comparison. The threshold was set at 5.13% and any module with a percentage of mitochondrial genes exceeding this threshold was classified as enriched.

Modules identified as enriched in mitochondrial genes were considered of particular interest for variant prioritization performed by VIOLA. These mitochondrial-enriched modules are likely to contain genes that play critical roles in mitochondrial function and are thus prioritized for further investigation.

#### Functional Enrichment Analysis

An enrichment analysis was performed using the EnrichR package in R^37^.

For this analysis, we queried nine distinct libraries, each offering insights into gene functions, phenotypes, and disease associations. In particular, we used the Gene Ontology (GO)^38, 39^ for Biological Process, Molecular Function, and Cellular Component (2023 version), to categorize genes based on their roles in biological systems, molecular activities, and subcellular locations. We also integrated the Human Phenotype Ontology (HPO)^40^ to explore links between genes and phenotypic abnormalities, as well as Rare Diseases GeneRIF and AutoRIF^41^ to investigate gene-disease relationships in the context of rare diseases. Additionally, we included the OMIM Disease library^42^, which integrates information from the Online Mendelian Inheritance in Man database, and Jensen DISEASES^43^, a resource offering curated associations between genes and diseases. We also incorporated ClinVar database^44^ (2019 version), which catalogues clinically relevant genetic variants and their roles in human health.

The results of the enrichment analysis were filtered to retain only statistically significant findings, defined as those with a p-value less than 0.05.

### VIOLA workflow

VIOLA integrates genomics, phenomics and transcriptomics to identify rare variants with peculiar combination of scores which could be potential candidates for diagnosis of very rare and particular MD. The workflow is composed of three main blocks that are described below (Figure 1).

**Figure 1:**
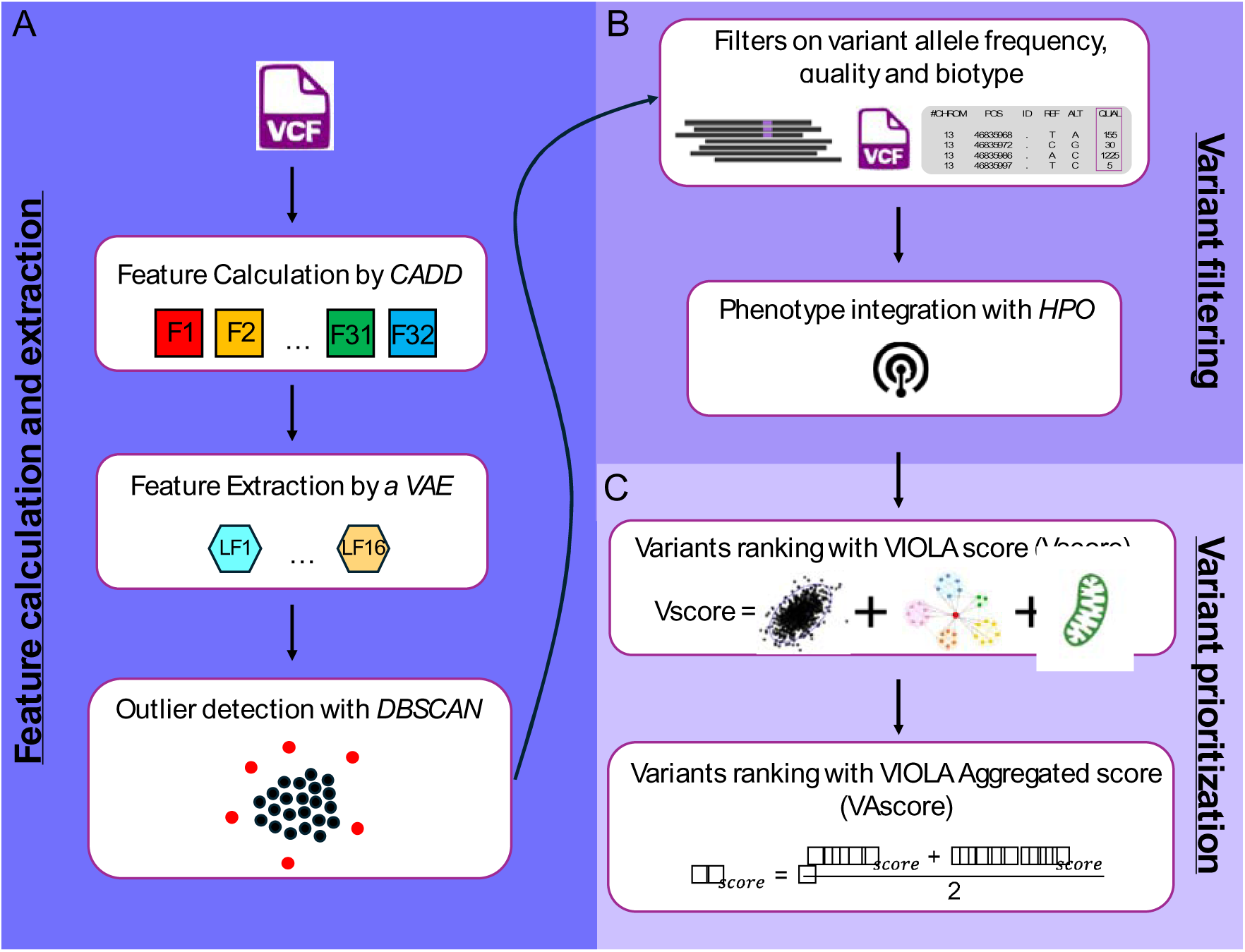
Overview of the VIOLA Pipeline. A) The VIOLA pipeline begins with feature calculation using CADD from a VCF file. These 32 features (F1, F2…) are then transformed using a Variational Autoencoder (VAE) into 16 latent features (LF1, LF2…), followed by outlier detection with the DBSCAN algorithm. B) The second stage of VIOLA focuses on filtering false positive variants. This is achieved by applying filters based on variant allele frequency (VAF), variant quality, and variant biotype. Next, the patient’s phenotype is integrated using the Human Phenotype Ontology (HPO), retaining only variants that could potentially explain the observed phenotype. C) In the final step, variants are ranked using different scoring methods. The first, the VIOLA score (Vscore), incorporates the Mahalanobis distance, transcriptomics data via a co-expression approach, and features related to mitochondrial diseases (MD). The second, the VIOLA Aggregated Score (VAscore), combines the Vscore with the Exomiser score, a state-of-the-art tool for variant prioritization.

#### Step A: Variant features calculation

To characterize genetic variants, we used the Combined Annotation-Dependent Depletion (CADD) tool^17^ which integrates multiple genomic features into one score to evaluate the deleteriousness of variants (Figure 1A).

Therefore using CADD, we calculated 32 features to describe variants, including six conservation scores from phastCons and PhyloP tools^45^, two pathogenicity scores from two functional impact predictors: SIFT^15^ and PolyPhen^16^ and two scores calculated by CADD^17^. We added eight epigenetic scores from ENCODE catalog^46^ and five splicing scores from MMSplice tool^47^. Finally, the nine remaining scores were calculated with BRAVO database^48^ to include the genomic environment of variants.

#### Step B: Techniques of dimensionality reduction

We used a variational autoencoder (VAE) to find complex non-linear combinations of variants features and reducing the dimensionality.

The VAE was built with Tensorflow^49^ and is composed of one hidden layer in the encoder of dimension 25 and the latent space which contains 16 latent features. Each layer of the encoder or decoder is composed of the dense layer followed by a batch normalization and the ELU activation function. To optimize the model, a custom loss function was implemented, combining the Mean Squared Logarithmic Error (MSLE) reconstruction loss and the Kullback–Leibler (KL) divergence to enforce latent space regularization.

We fed the VAE with the variants scores calculated in step A to obtain 16 new features that are combinations of the 32 original variants scores (Figure 1A).

#### Step C: Outlier detection

Since we reasoned that the responsible variant for MD suspicions should have a very peculiar combination of parameters, this should be seen as an outlier in the space of variants parameters. We employed the Density-Based Spatial Clustering of Applications with Noise (DBSCAN) algorithm as a first step toward the identification of the outliers’ variables. DBSCAN is an unsupervised machine learning algorithm used for clustering spatial data based on density. Unlike traditional clustering methods, DBSCAN identifies clusters as dense regions of points separated by areas of lower point density, which allows it to effectively handle datasets with noise and outliers. DBSCAN requires two parameters: the maximum distance two points belonging to the same cluster (epsilon, ε) and the minimum number of points required to form a cluster (minPts).

For datasets with more than two dimensions, it is recommended to set the *minPts* parameter to two times the number of dimensions, therefore we settled to 32^50^.

To determine an appropriate ε, we used a data-driven approach based on the k-nearest neighbors (k-NN) method^51^. Specifically, for each point, we computed the average distance to its k-nearest neighbors, where k was set to √N (with *N* being the number of variants). These distances were then sorted in ascending order, and the final ε value was chosen as the mean of the first *n* distances (where *n* was empirically set to 1500). Based on those parameters, DBSCAN divides the variants in two groups: the outliers and the non-outliers. We retrieve only the variants classified as outliers to continue the pipeline (Figure 1A).

#### Step D: Variant filtering

We apply a series of filters to variants retained from the previous step (Figure 1B). First, we removed all low-quality variants by excluding those with a quality score below 50. This threshold was chosen to minimize false positives, as variants with lower quality scores are often the result of sequencing errors or low-confidence base calls. Next, since our study is based on WES data, we focused exclusively on variants located in protein-coding regions or which have an impact on a protein-coding gene. While WES primarily targets the exonic regions of the genome where the majority of known pathogenic variants are found, we also included intronic variants if they were predicted to affect the function or regulation of a protein-coding gene. By filtering for “protein-coding” variants, we reduced the data to include only those variants that are likely to have functional effects on gene products, such as nonsynonymous variants, splice-site alterations, and frameshift mutations. Finally, we applied a final filter based on variant allele frequency (VAF), which measures the proportion of variant alleles at a genomic locus. For homozygous variants, the VAF must be greater than 0.8, ensuring that the variant is present in most sequencing reads at that position. Conversely, for heterozygous variants, the VAF should fall between 0.3 and 0.7, reflecting a balanced presence of both reference and alternative alleles. Variants that do not meet these criteria are filtered out as they may result from sequencing errors, mapping biases, or other technical artifacts.

#### Step E: Phenomics integration

To integrate the phenotype of the patient, we used the Human Phenotype Ontology (HPO)^40^. HPO is a database that provides a standardized vocabulary for describing phenotypic abnormalities associated with human diseases.

To quantify the similarity between two phenotypic terms, we calculated the Jaccard Index for the sets of ancestors associated with the two HPO terms. We defined the Jaccard index of HPO terms as:

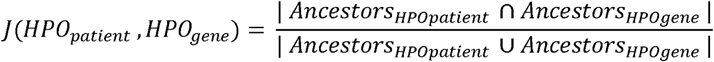

The index gives values ranging from 0 to 1, where a value of 0 indicates that there are no shared elements between the sets (no similarity), and a value of 1 indicates that the sets are identical (complete similarity). Therefore, a high Jaccard index indicates greater similarity between the phenotypic features represented by the terms, while a low Jaccard index represents a small overlap (Figure 1B).

#### Step F: filtering variants based on the Mahalanobis distance

To identify variants that significantly deviate from the distribution of CADD scores, we computed the Mahalanobis distance^52^, which measures the distance between a point and the mean of a multivariate distribution while considering correlations between variables. This approach allows for the detection of outliers based on multiple features simultaneously. Thus, we first selected the 32 scores extracted in step A, then replaced missing values with the median of the corresponding score and finally computed Mahalanobis distance with the function *mahalanobis* from stats package^53^.

To detect significantly distant variants, we calculated the p-value associated with the Mahalanobis distance using the Χ^2^ distribution. Variants with p-value < 0.001 were considered outliers.

#### Step G: Transcriptomics data integration and definition of a first ranking score for variants: the VIOLA Score (Vscore)

We defined a first score which includes multiple parameters to provide a first score to rank variants accordingly to the disease (Figure 1C). First, we included transcriptomics information leveraging MitoBook using a binary score. This score indicates whether a variant is associated with a gene belonging to a module enriched in mitochondrial genes, identified by co-expression analysis using WGCNA. Variants linked to genes belonging to such modules are assigned a score of +1, while those outside these modules are assigned a score of -1. Details of this process are described in the Materials and methods section *MitoBook: a catalog of co-expressed genes modules in MD context*.

A second parameter used considers whether a variant is unique and specific to a patient, with the exception of cousins for whom we examined common variants not present in the rest of the cohort. Variants meeting this criterion are assigned a score of +1, while non-unique variants are given a score of 0. Prior knowledge regarding genes involved in MD was also included. Referring to Stenton and Prokisch^11^, 413 distinct genes are associated with MD. Variants in these known genes are assigned a score of +1, as they are more likely to be implicated in the disease. Variants in genes not yet linked to MD are assigned a score of 0.

Finally, although the Mahalanobis distance was already used to filter variants accordingly to the p-value associated to the measure, we also decided to include as additional term in the Vscore to consider the degree of significance provided by the statistical test.

To balance the contributions of the four components of the final Vscore, we assigned specific weights to each score. The Mahalanobis distance, transcriptomics, and uniqueness features were each assigned equal weights of 0.5, reflecting their comparable importance. In contrast, the score for known MD genes was assigned a lower weight of 0.01. This lower weighting accounts for the heterogeneity of MD and the incompleteness of the current MD gene catalog.

The Vscore is calculated as the weighted sum of the Mahalanobis distance and the scores for transcriptomics, uniqueness, and known MD genes:

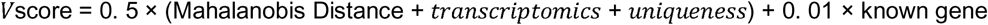

Finally, the Vscore undergoes a normalization process using *MinMax* normalization to ensure that all scores fall within a standardized range, facilitating comparison across different variants. Once normalized, the variants are ranked based on their adjusted Vscore. Then, we apply a filtering step to retain only the most relevant variants by selecting those with a normalized Vscore greater than the third quartile (Q3), allowing us to focus on the top-ranking variants.

#### Step H: Definition of the Autosomal Recessive rank (ARrank)

Since most of the genes known to be involve in MD are AR inherited, we created a rank specific for AR inheritance in which we incorporated the genotype of the patient called ARrank.

Variants are filtered according to their genotype. Only homozygous and compound heterozygous variants are compatible with AR mode of transmission, so we have kept all genes with homozygous variants and genes with at least two heterozygous variants.

The rank of the remaining variants is then recalculated still using the Vscore.

#### Step I: Definition of the VIOLA Aggregated score (VAscore)

To create the VAscore, we calculated the arithmetic mean of the Vscore and the Exomiser score for each variant (Figure 1C). The Exomiser score ranges from 0 to 1 so each score contributes equally to the final VAscore as they have the same range.

#### Step J: Output of VIOLA

To improve the accessibility of our results, we have opted for a more interactive and user-friendly approach by providing the results in the form of an HTML file rather than as static tables, which can often be difficult to interpret. Interactive HTML tables bring together the most important information in a simplified format. These tables include key columns and scores, such as SIFT^15^ and PolyPhen^16^ values, MitoCarta database^36^ and variant frequencies from gnomAD^31^ and 1000G^32^. This gives users rapid access to the most relevant data.

Databases such as dbSNP^30^, gnomAD^31^, OMIM^42^, GeneCards^54^, Genotype Tissue Expression (GTEx)^55^ and Ensembl^56^ are included to provide an overview of the gene or variant in question. By clicking on the links provided, users are directed to the relevant database entry in a new tab, offering immediate access to detailed information on the variant or gene in question.

### Implementation

VOLA is implemented in Python and R. The only two steps which are done in python are the use of the VAE and DBSCAN to find outliers. The rest of the pipeline is done in R.

### Benchmark of VIOLA

#### Dimensionality reduction method

To test if the association of a technique of dimensionality reduction and DBSCAN is needed to retrieve the responsible variants among the outliers we tested different configurations. First tried without any dimensionality reduction technique, therefore applying directly DBSCAN on CADD scores.

Then, we tested other dimensionality reduction method instead of the VAE: Principal Component Analysis (PCA) and t-Distributed Stochastic Neighbor Embedding (t-SNE).

##### Principal Component Analysis (PCA)

We implemented the PCA^57^ using the scikit-learn library^58^. First, PCA was performed without specifying the number of components, allowing the computation of the cumulative explained variance across all principal components. The optimal number of principal components was determined by selecting the smallest number of components that explained at least 95% of the total variance.

Once the optimal number of components was identified, we refitted PCA using this selected number of components and transformed the dataset accordingly. The transformed features were stored in a new dataset, where each column corresponds to a principal component.

##### T-Distributed Stochastic Neighbor Embedding (t-SNE)

We applied t-SNE^59^ which is a non-linear dimensionality reduction technique designed for exploring complex structures in high-dimensional data. We used the scikit-learn implementation of t-SNE^58^ to project the feature space into two dimensions.

#### Shuffle analysis

To validate the basic assumption underlying our new model, namely that the combination of CADD scores is an essential element for pipeline success, we conducted a shuffling experiment. This experiment was designed to test the extent to which the structure and accuracy of CADD scores influence overall pipeline performance.

Our approach focused on the four patients in the cohort for whom the causal variant had already been identified. For each patient, we took the CADD scores of the variants and applied a shuffling strategy to disrupt their original associations while preserving their statistical distribution. Specifically, shuffling was performed on a column-by-column basis, ensuring that the overall distribution of CADD scores remained identical to that of the original dataset, but with random assignments to individual variants. This method enabled us to isolate the impact of the original, unshuffled CADD scores on the downstream stages of the pipeline.

To ensure statistical robustness, we repeated the shuffling process 100 times, generating 100 distinct datasets with shuffled CADD scores. For each shuffled dataset, we ran the entire pipeline, starting from the application of the VAE to calculation of Vscore.

#### Exomiser

We ran Exomiser using a standardized pipeline that automates the configuration process for each patient. Since Exomiser requires a configuration file specifying input VCF files, HPO terms, and output paths, we developed a Python script to dynamically generate patient-specific configuration files. This script reads a table containing patient IDs, corresponding VCF file paths, and associated HPO terms, then modifies a template Exomiser configuration file accordingly. To efficiently process multiple patients, we generated a batch-commands file that lists all configuration file paths, enabling the execution of Exomiser in batch mode on a high-performance computing server. This automated approach ensured reproducibility and scalability in variant prioritization across the cohort.

## Results

### Overall study design

VIOLA was performed on sequencing data from a cohort of patients. All patients were referred to the reference center at Nice University Hospital for suspected mitochondrial diseases (MD). Detailed characteristics of this cohort are presented in Table 1. The cohort includes 20 patients of which 13 males and 7 females. Two of the patients in the cohort are cousins (patient 3 and patient 17); patients range in age from 24 hours to 50 years old.

As part of the diagnostic workflow, all patients underwent blood testing for Whole-Exome Sequencing (WES) and skin biopsies to extract fibroblasts for RNA sequencing.

Blood is commonly used for WES because it is easily accessible and minimally invasive for identifying germline variants. On the other hand, for RNA-seq, the choice of the tissue is very important since the gene expression can vary significantly between tissues. Fibroblasts were chosen because as a compromise between tissue accessibility, easy to culture *in vitro* and representativeness of the major impacted tissue of MD (muscle). In addition, many genes with mitochondrial function have been shown to be expressed in fibroblasts^60^. A comparison was made between gene expression in skin fibroblasts and whole blood. The results showed that gene expression in fibroblasts is more consistent and has less variability compared to whole blood, making fibroblast RNA-seq preferable for detecting biologically and clinically relevant gene expression differences^61^.

Among the 20 patients, the causal variant responsible for the suspected disease was identified in 4 cases, providing a confirmed genetic diagnosis. However, for the remaining 16 patients, the classical diagnostic process remains inconclusive, leaving these cases in a diagnostic stalemate, which motivated this work.

### VIOLA: a new model to prioritize genetic variants

To develop a tool to perform variants prioritization for patients with suspected MD when the responsible variant is not present in common genes involved in MD, we performed an extensive benchmark and selected the best performing tools. Our hypothesis is that since the disease-responsible variant(s) is (are) individual-specific and very rare, therefore it (they) should have a unique combination of properties different from the other individual’s variants. Hence, we assume that the putative disease variants for MD suspicions are outliers of each individual variants’ distribution. VIOLA consists in three main blocks: feature calculation and feature extraction; variants filtering; variants prioritization.

Regarding the features calculation to represent different characteristics of variants, we provide a module which extracts 32 scores calculated by CADD spanning pathogenicity to splicing to characterize all types of variants and not only coding variants. To extract the non-linear combination of variants characteristics, we employed a VAE which provide in the latent space a compressed representation of complex patterns hidden in the input data.

Then, since the assumption is that the responsible variant is an outlier, we used a density-based clustering technique called DBSCAN directly on the latent space to retrieve the outliers’ variants.

Furthermore, different filters on biotype, variant quality and variant allele frequency and the integration of the patient phenotype with HPO terms are used to reduce the number of potential responsible variants.

The last steps consist in the calculation of two scores for variant prioritization. The Vscore is based on Mahalanobis distance metric and the combination of several features related to MD and the VAscore which is the combination of the Vscore and Exomiser score.

VIOLA generates both a tabulated file and an interactive HTML file to improve accessibility for non-bioinformatician users. The HTML file allows easy navigation and is directly linked to widely used genetic databases.

We applied the pipeline on collections of rare variants from 20 patients with suspected MD including 4 with known causal variants.

### Benchmark of feature calculation and feature extraction

To validate our approach and assess the robustness of our pipeline, we designed and tested several scenarios using data from the four patients for whom the causative variants are already known. These scenarios were designed to assess, firstly, the need for a dimensionality reduction technique and, secondly, which technique is the most effective. All the results are reported in Table III.

**Table III:** Summary of benchmarking results for dimensionality reduction methods. In the table, results of DBSCAN clustering applied to genetic variants, with and without dimensionality reduction techniques are reported. The table includes the method used for dimensionality reduction such as variational autoencoders (VAE), principal component analysis (PCA), t-Distributed Stochastic Neighbor Embedding (t-SNE), or none, the number of selected features, the distribution of variants between the two DBSCAN classes (outlier and non-outlier), and the classification of the known disease-causing variant.

| Method | Patient | Number of features | Nb_non-outlier-variant | Nb_outlier_variant | Cluster_of_responsible_variant |
| --- | --- | --- | --- | --- | --- |
| VAE | 3 | 16 | 40830 | 50128 | outlier |
| VAE | 4 | 16 | 78308 | 93011 | outlier |
| VAE | 9 | 16 | 5674 | 5959 | outlier |
| VAE | 17 | 16 | 37652 | 46767 | outlier |
| PCA | 3 | 13 | 0 | 90958 | outlier |
| PCA | 4 | 11 | 44 | 171275 | outlier |
| PCA | 9 | 14 | 0 | 11633 | outlier |
| PCA | 17 | 13 | 0 | 84419 | outlier |
| TSNE | 3 | 2 | 90958 | 0 | Non-outlier |
| TSNE | 4 | 2 | 171319 | 0 | Non-outlier |
| TSNE | 9 | 2 | 11633 | 0 | Non-outlier |
| TSNE | 17 | 2 | 84419 | 0 | Non-outlier |
| No | 3 | 32 | 0 | 90958 | outlier |
| No | 4 | 32 | 0 | 171319 | outlier |
| No | 9 | 32 | 0 | 11633 | outlier |
| No | 17 | 32 | 0 | 84419 | outlier |

In the first scenario, we tested the pipeline without using the VAE (Figure 2A). Hence, we applied the DBSCAN clustering algorithm directly to the 32 scores calculated by CADD. However, this approach proved ineffective, as all variants were assigned to the outlier group by DBSCAN. This result demonstrates that clustering the raw CADD scores without dimensionality reduction do not provide meaningful information for prioritizing potentially pathogenic variants in the context of MD.

**Figure 2:**
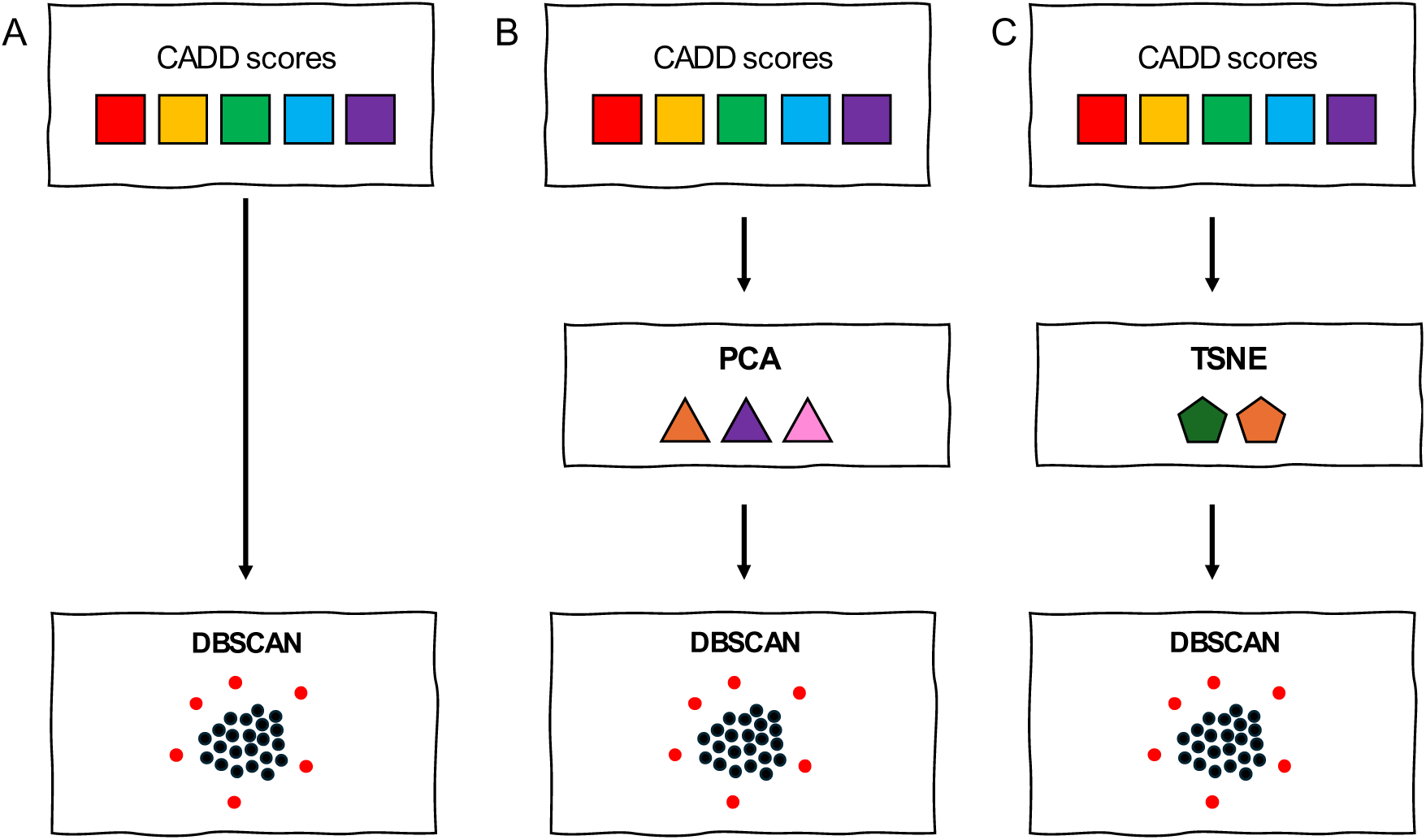
Comparative analysis of dimensionality reduction methods across different scenarios. In the first scenario, we applied DBSCAN directly to the CADD score without using a variational autoencoder (VAE) (A). In the other two scenarios, we replaced the VAE with either PCA (B) or t-SNE (C) before applying DBSCAN, following the same approach as in the original pipeline.

To further explore alternative approaches for dimensionality reduction, we set up two additional scenarios. Therefore, we replaced the VAE with two widely used dimensionality reduction methods: PCA (Figure 2B) and t-SNE (Figure 2C).

By feeding the PCA algorithm with the 32 scores, we extracted between 11 and 13 features for the four patients with known causal variant, yielding between 11633 and 90958 variants classified as outliers by DBSCAN, indicating that PCA failed to adequately capture the critical features necessary for effective clustering (Table III). PCA’s linear nature may not have been sufficient for modeling the complex relationships in the data. Similarly, when t-SNE was used as feature extraction technique, the responsible variants were not classified as outliers. t-SNE failed to differentiate between the causative variants and the majority of benign variants.

Those results rule out PCA or t-SNE as alternative tools to VAE for feature extraction to be included in VIOLA.

The application of this step to our cohort filtered out 45% of variants (Figure 3, Table IV, Supplementary Table I).

**Figure 3:**
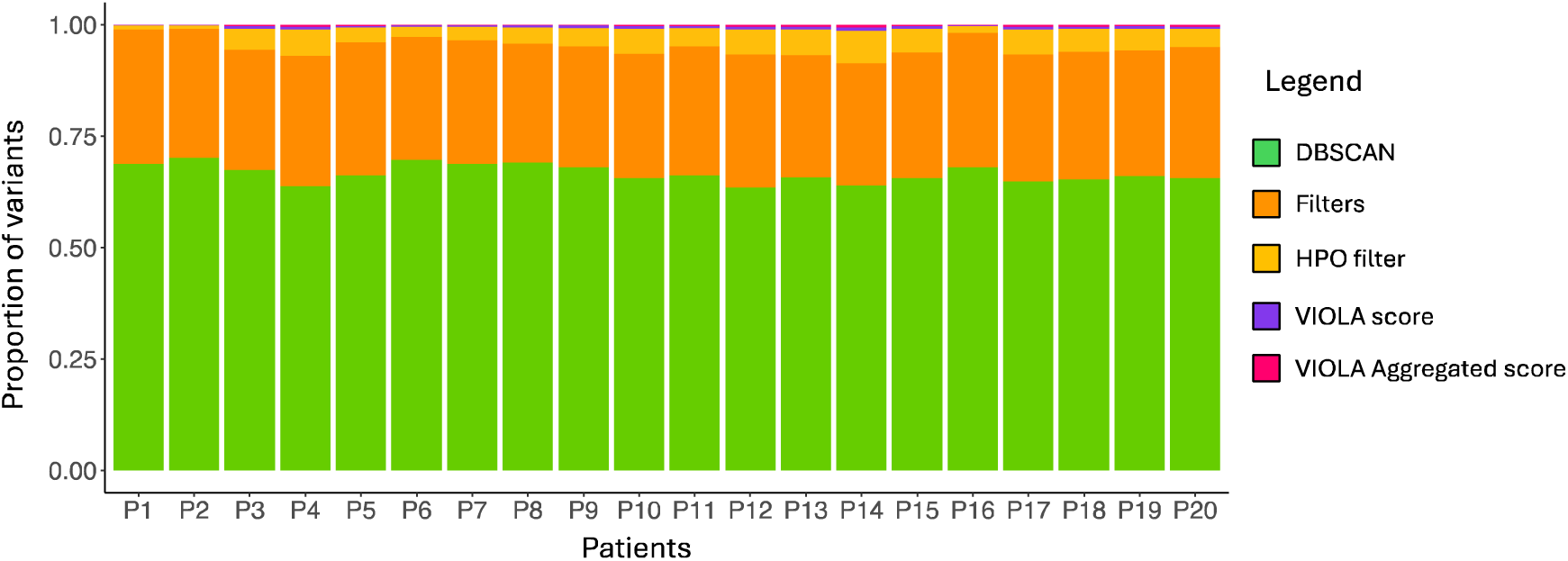
Proportion of retained variants at each step of the VIOLA pipeline across patients. The stacked barplot represents the percentage of retained variants at each step of the VIOLA pipeline for each patient in the cohort. The x-axis corresponds to individual patients, while the y-axis shows the percentage of variants remaining after each step: outlier detection (green), filters on variant allele frequency (VAF), variant quality and variant biotype (orange), phenotype integration (yellow), VIOLA rank (Vrank) (purple) and VIOLA Aggregated rank (VArank) (pink).

**Table IV:** Percentage of filtered variants at each step of the VIOLA pipeline for all patients. The percentage of filtered variants at each step of the VIOLA pipeline is reported in the table from the DBSCAN step to the ranking with the Autosomal Recessive rank (ARrank) for the 20 patients of the cohort. Abbreviations for steps are DBSCAN: Density-Based Spatial Clustering of Applications with Noise, HPO: Human Phenotype Ontology, Vrank: VIOLA rank, ARrank: Autosomal Recessive rank.

| Patient | DBSCAN | Filters | HPO | Vrank | ARrank |
| --- | --- | --- | --- | --- | --- |
| 1 | 44 | 56 | 97 | 89 | 39 |
| 2 | 46 | 59 | 97 | 88 | 24 |
| 3 | 45 | 60 | 82 | 89 | 27 |
| 4 | 46 | 54 | 80 | 90 | 16 |
| 5 | 45 | 55 | 89 | 89 | 17 |
| 6 | 47 | 60 | 92 | 85 | 69 |
| 7 | 47 | 60 | 89 | 86 | 65 |
| 8 | 47 | 61 | 87 | 86 | 67 |
| 9 | 49 | 60 | 85 | 85 | 62 |
| 10 | 44 | 57 | 80 | 89 | 35 |
| 11 | 44 | 56 | 86 | 89 | 23 |
| 12 | 44 | 53 | 81 | 89 | 29 |
| 13 | 44 | 58 | 79 | 89 | 30 |
| 14 | 45 | 57 | 73 | 89 | 25 |
| 15 | 44 | 57 | 81 | 89 | 29 |
| 16 | 45 | 56 | 95 | 88 | 30 |
| 17 | 45 | 56 | 80 | 89 | 31 |
| 18 | 44 | 56 | 82 | 89 | 25 |
| 19 | 44 | 57 | 83 | 89 | 27 |
| 20 | 45 | 55 | 86 | 88 | 16 |

### The integration of phenomics considerably reduces the number of variants classified as potential responsible for the disease

After identifying the outlier group of variants using our pipeline, we applied a series of additional filters to refine the prioritized list of variants. These filters focused on three key criteria: the quality of the variants, the biotype of the genes affected by these variants and the VAF. Variants with low-quality scores were excluded to ensure the reliability of the analysis. Additionally, we restricted our analysis to variants located in protein-coding regions or other relevant biotypes known to play significant roles in disease mechanisms. The last filter concerned the VAF to remove variants that were inconsistent in terms of variant genotype and allele frequency. For more details on filters, see from Materials and methods. This first step resulted in a significant reduction in the number of variants to be screened, with an average of 57% of variants being eliminated at this stage (Figure 3, Table IV, Supplementary Table I).

The next step focused on the identification of the variants relevant to the patient’s phenotype. Therefore, we integrated phenotypic information from HPO database. Clinicians have annotated the patients in the study with their correspondent HPO term(s) that exhaustively described the observed clinical phenotype. The challenge was to determine whether a given variant was linked to a gene associated with phenotypic features similar to those of the patient.

Therefore, we developed a novel approach based on the Jaccard index computation between the ancestors of HPO terms associated with the genes bearing variants and the HPO terms representing the patient’s phenotype. This approach was iterated for all possible pairwise combinations of HPO terms associated with the variants and those of the patient. After calculating the Jaccard index values, we arbitrarily established a threshold of 0.80 to retain only those variants associated with genes whose HPO terms showed significant similarity to the patient’s phenotype. By applying this threshold, we ensured that the retained variants were likely to be functionally relevant to the clinical features observed in the patient. This is the most stringent step before the variants are ranked. On average, we excluded 85% of variants (Table IV, Supplementary Table I).

This score enables to include phenomics considerations into the selective process to identify the individuals’ responsible variants.

### Towards a strategy to include transcriptomics using co-expression network analysis

To explore the relationships between genes and mitochondrial functions, we developed MitoBook a comprehensive catalog of gene co-expression patterns in patients with suspected MD. Briefly, we applied a co-expression analysis to the transcriptomics data from the 20 patients from the cohort which we dispose. We applied WGCNA^35^ to identify 12 distinct co-expression modules. These modules vary in size, ranging from 64 to 11,617 genes. Notably, the largest module, containing 11,617 genes, comprised almost half of the genes in the input expression matrix. Such a large module, lacking specificity, likely represents a background or generic expression pattern, it was excluded from further analyses to focus on modules with greater biological relevance. Similarly, the smallest module, consisting of only 64 genes, was also excluded.

Then, we investigated whether the 10 remaining modules were enriched in mitochondrial genes. To do this, we calculated a “propensity score” using the MitoCarta database^36^ (see Materiel and Methods, *Enrichment in mitochondrial genes*). This analysis enabled us to identify eight modules that show a significant enrichment in mitochondrial genes, ranging from 5.36 to 16.54%, indicating their propensity to be relevant in mitochondrial biology (Figure 4A). The analysis was extended to additional nine databases containing information about the genes functions in mitochondrial functioning. The results are presented in the form of a heatmap showing, in rows, the different mitochondrial pathways we found, and, in columns, the modules found by WGCNA. The color scale represents the -log10 of the pvalue (Figure 4B). The strongest enrichments were observed in “greenyellow” module which consists of 184 genes, 14.13% of which are mitochondrial genes. This module showed a significant overrepresentation of genes involved in mitochondrial ATP synthesis coupled to electron transport, as well as genes associated with mitochondrial respiratory chain complex I and NADH dehydrogenase activity. This module groups together genes that play a crucial role in mitochondrial energy production and electron transport processes.

**Figure 4:**
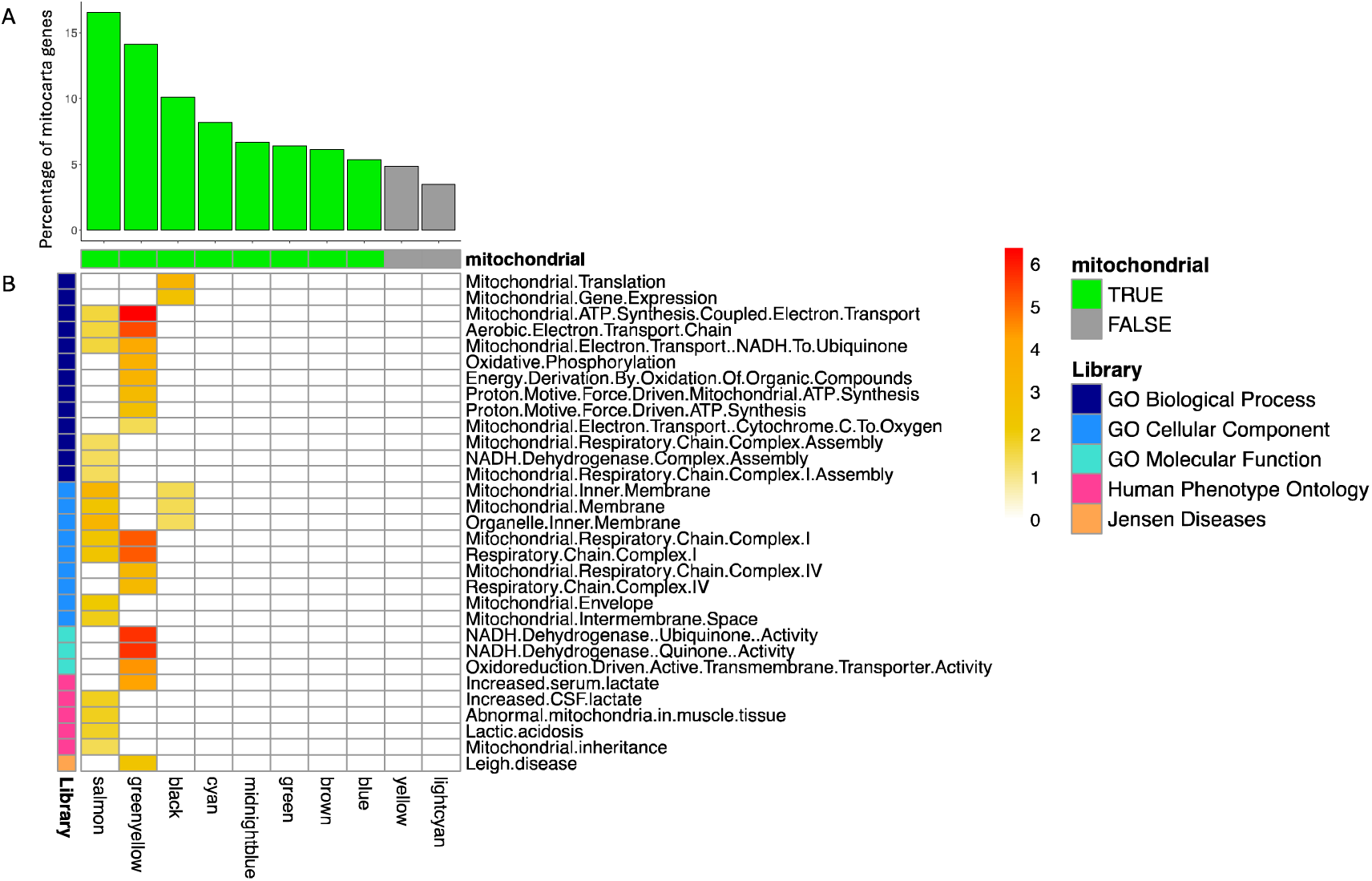
Mitochondrial gene proportion and pathway enrichment in co-expression modules. The barplot (A) shows the percentage of mitochondrial genes in each module identified by co-expression analysis. Modules highlighted in green are enriched in mitochondrial genes, while those in gray are not. The heatmap (B) presents the results of functional enrichment analysis in five different ontologies: Gene Ontology (GO) Biological Process (dark blue), GO Cellular Component (turquoise), GO Molecular Function (cyan), Human Phenotype Ontology (pink), and Jensen Diseases (salmon). The x-axis represents the modules, while the y-axis lists the mitochondrial pathways identified. The color scale indicates the strength of enrichment, with more intense colors signifying stronger associations.

Overall, three modules with the highest propensity score from MitoCarta also achieved the highest score for the five other databases: GO Biological Process, GO Cellular Component, GO Molecular Function, HPO and Jensen Diseases. This convergence of evidence across multiple resources underlines the functional relevance of these three modules and highlights their potential importance in understanding mitochondrial processes and their links to patient phenotypes.

These results conducted to the definition of a score based on the guilty-by-association principle. Therefore, genes present in a module with high propensity score (> 5.13), were assigned with a MitoBook score of 1 otherwise of -1.

The MitoBook score allows to consider an additional omics layer, the transcriptomic to be added to the overall estimation of the degree of pathogenicity.

### Evaluating the ranking efficacy of Vscore

To summarize all the calculated scores in one unique value to be used to prioritize potential pathogenic variants for MD suspicions, we first defined a novel comprehensive score, the VIOLA score (Vscore).

The Vscore incorporates multiple features, such as the Mahalanobis distance, the MitoBook score, and additional metrics related to mitochondrial diseases, to rank variants according to their likelihood of being pathogenic.

We chose Mahalanobis distance because it is a spatial measure and allows us to filter variants and detect outliers that significantly differ from the distribution of selected features. We added Mitobook score to integrate transcriptomics data to better prioritize variants into genes that show similar expression profile to known mitochondrial genes. We also considered a score that reflects the uniqueness of the variant, since a variant that occurs more than once in our cohort (except for related patients) is less likely to be the responsible variant. Finally, we incorporated a score based on genes already known to be involved in MD to add prior knowledge about the disease.

From the Vscore, we created two ranks, the Vrank which is general and the ARrank which includes genotype-specific filters. Exomiser score considers variant pathogenicity, frequency and the patient phenotype.

We developed two separate ranks because since most of the genes already involved in MD have an autosomal recessive (AR) inheritance, we wanted to give importance to genes compatible with AR inheritance by creating the ARrank while keeping a general rank with all the variants prioritized by VIOLA regardless of transmission mode (Vrank).

We evaluated the efficacy in ranking for both ranks of the Vscore on the four patients for whom the causal variants were already known, comparing the ranking obtained with Exomiser, the state-of-the-art tool to perform variant prioritization. The results are summarized in table V.

**Table V:** Ranks of responsible variants for diagnosed patients across different prioritization scores. The table presents the ranking positions of the causal variants for the four diagnosed patients in the cohort, as determined by different variant prioritization scores. The rankings were obtained using VIOLA rank (Vrank), Autosomal Recessive rank (ARrank), VIOLA Aggregated score (VAscore), and Exomiser.

| Patient | Vrank | ARrank | Rank with VAscore | Rank with Exomiser score |
| --- | --- | --- | --- | --- |
| 3 | 1 | 1 | 1 | 9 |
| 4 | 7 | 3 | 3 | 18 |
| 9 | 5 | 4 | 1 | 19 |
| 17 | 1 | 1 | 1 | 17 |

For patients 3 and 17, the causative variant was ranked at the first place using both VIOLA ranks, it was ranked respectively at 9^th^ and 17^th^ positions for Exomiser (Figure 5A and 5B). This indicates that the Vscore was able to identify the pathogenic variant as the most likely candidate, confirming its reliability in variant ranking.

**Figure 5:**
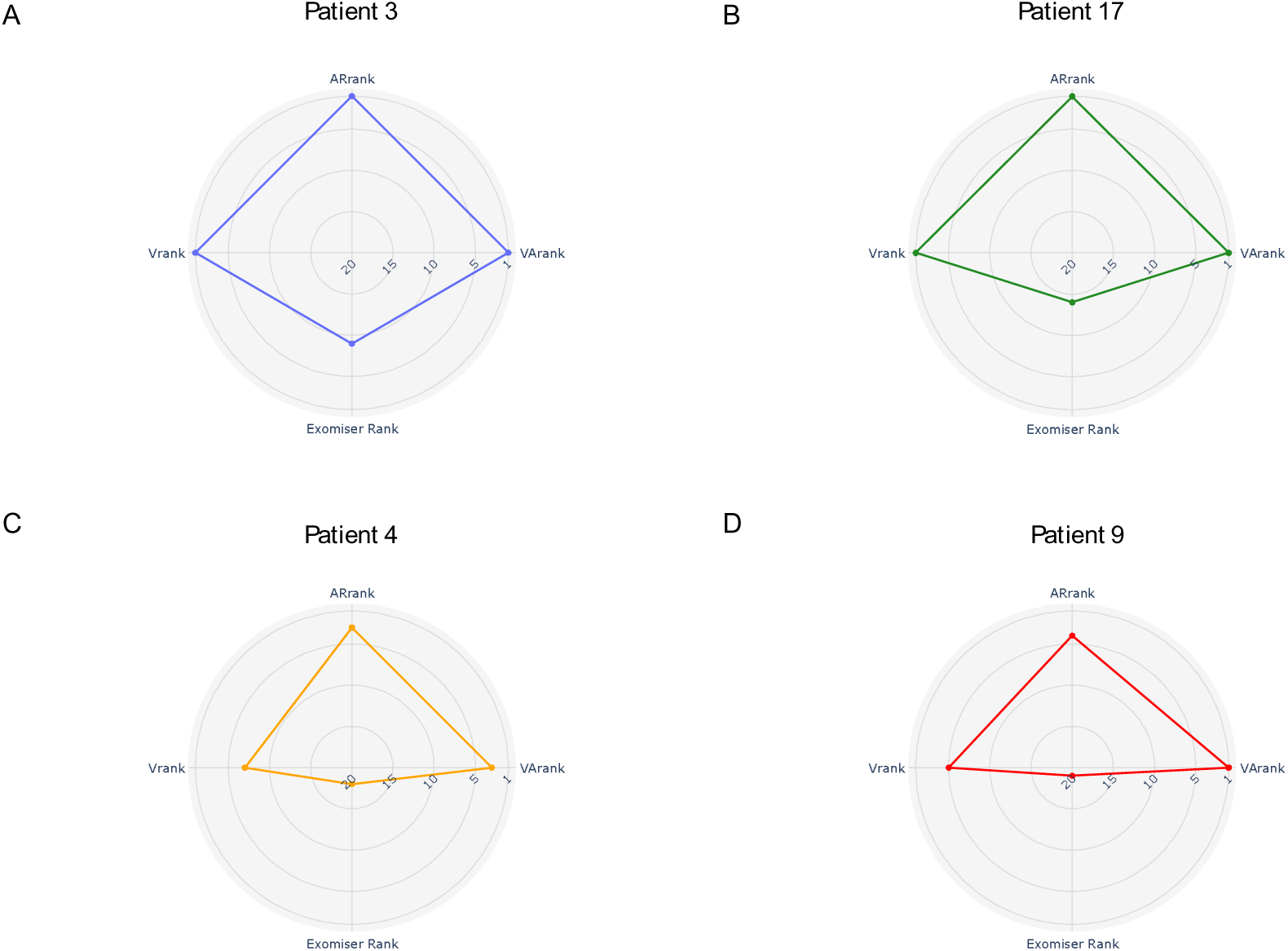
Comparison of responsible variant ranks using different scoring methods for diagnosed patients. Radarplots illustrate the ranks of the responsible variants obtained with the VIOLA rank (Vrank), the Autosomal Recessive rank (ARrank), the VIOLA Aggregated rank (VArank) and the Exomiser rank for patient 3 (A), patient 17 (B), patient 4 (C) and patient 9 (D).

For patient 4, the Vscore reported the responsible variant in the 7^th^ position for the Vrank while it was 18^th^ with Exomiser. However, if the ARrank is used, the rank improved significantly, placing the variant in the 3^rd^ position (Figure 5C).

Similarly, for patient 9, the responsible variant achieved a rank of 5^th^ w^33^hen using the Vrank. With the ARrank, the rank improved further, elevating the variant to the 4^th^ position while with Exomiser, the variant was ranked 19^th^ (Figure 5D).

The results highlight the added value of integrating genotype-specific data to refine the prioritization process, particularly in cases where the responsible variant may not stand out as the top candidate based solely on other features.

### VIOLA Aggregated score results

Based on the previous results, we reasoned to define an integrated score considering both Exomiser score and Vscore into a single score, the VIOLA Aggregated Score (VAscore).

For patients 3 and 17, the VAscore reported the responsible variant at the top of the ranking as for the Vrank only, suggesting that the Exomiser score is not influencing VIOLA results. Similarly, for patient 4, the responsible variant is still in 3^rd^ position, achieving the same rank than with ARrank.

The greatest improvement is observed for the responsible variant of patient 9. Using only the Vrank, the responsible variant is ranked in 5^th^ position, while Exomiser’s rank is 19^th^ position. However, with VAscore, the variant is upgraded to the first position, underlining the significant advantage of combining these scores.

These results demonstrate the value of combining Exomiser and VIOLA scores into a single score. By exploiting the unique strengths of each tool, the VAscore offers a more robust and accurate ranking system, improving the identification of pathogenic variants in complex rare disease cases. This integrated approach represents a significant advance in variant prioritization strategies, ensuring greater confidence in the identification of pathogenic variants.

Overall, the prioritization of responsible variants was compared with the rankings obtained by each tool independently. Importantly, the VAscore does not compromise the accuracy of VIOLA results because the Vscore rankings are not changing when the additional scores in VAscore yield lowest values.

### Validation by conducting shuffle analysis

To validate the VIOLA pipeline, since we do not dispose of another cohort, we performed a shuffled analysis on the feature values associated to each variant as the input of VIOLA. We designed an approach to randomize the feature values to assess the importance of these scores for the ranking of the causal variants. Using data from the four patients with known causal variants, we shuffled the features values 100 times while preserving their original distribution. After each shuffle, we applied the entire VIOLA pipeline.

At the DBSCAN clustering level, an interesting pattern emerged: the clustering of variants in the input files remained consistent with the results obtained using the original CADD scores. However, a significant effect was observed when calculating Vscore and Vrank. In 28% to 69% of the shuffled analyses, depending on the patient, the causal variant was filtered out and therefore not classified by VIOLA. For the remaining analyses where the causal variant was retained, we examined its rank distribution to assess the impact of shuffling, and all the results are summarized in table VI.

**Table VI:**
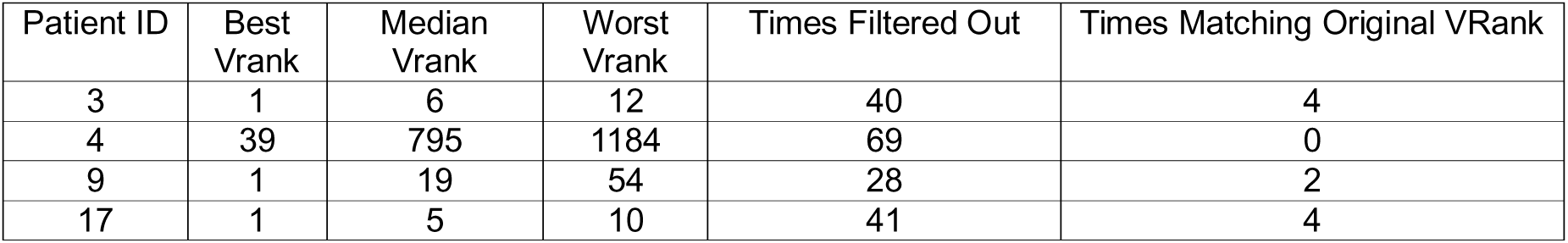
Summary of VIOLA Rank after CADD scores shuffling. This table summarizes the results of the CADD scores shuffle analysis performed on the four diagnosed patients. For each patient, the VIOLA pipeline was run 100 times with shuffled CADD scores. The table reports the best, median, and worst VIOLA rank (Vrank) obtained for the responsible variant, the number of times the variant was filtered out before ranking, and the number of times the original Vrank was recovered.

For patients 3 and 17, shuffling seems to have a less significant effect on causal variant ranking than expected. The Vrank ranged from 1 to 12 for patient 3 and from 1 to 10 for patient 17. The ranking obtained for the responsible variants from the original variants parameters was preserved in only four out of 100 of the shuffled analyses for both patients.

For patients 4 and 9 the shuffling impacted dramatically the ranking. For patient 9, the causal variant reached a maximum Vrank of 54 and a median of 19 and the original rank was only reproduced twice out of 72, while the highest Vrank of causal variant of patient 4 was 1184.

These results underline the crucial importance of variants parameters and their combination in our pipeline. The shuffled analyses reinforced the ability of VIOLA to accurately prioritize and identify the causal variant. This underlines that the pipeline we have developed is robust to noise and the underlined combination of parameters from original scores are fundamental for the stability and accuracy of the rankings.

### Results on the other patients of the cohort

We extended the application of VIOLA to the undiagnosed patients of our cohort. In absence of the knowledge about the responsible variants, we compared the performances of VIOLA and Exomiser regarding the prioritization of variants in mitochondrial-related genes. Therefore, we compared the proportion of variants located in genes listed in the MitoCarta database for both tools. Across the 20 patients in the cohort, VIOLA consistently identified a higher proportion of variants related to mitochondrial genes than Exomiser (Figure 6), suggesting that VIOLA is more specifically tailored to prioritize variants that are functionally relevant to mitochondrial biology. By focusing on features and pathways associated with mitochondrial function, VIOLA can better capture disease-relevant variants that may be overlooked by more general tools such as Exomiser.

**Figure 6:**
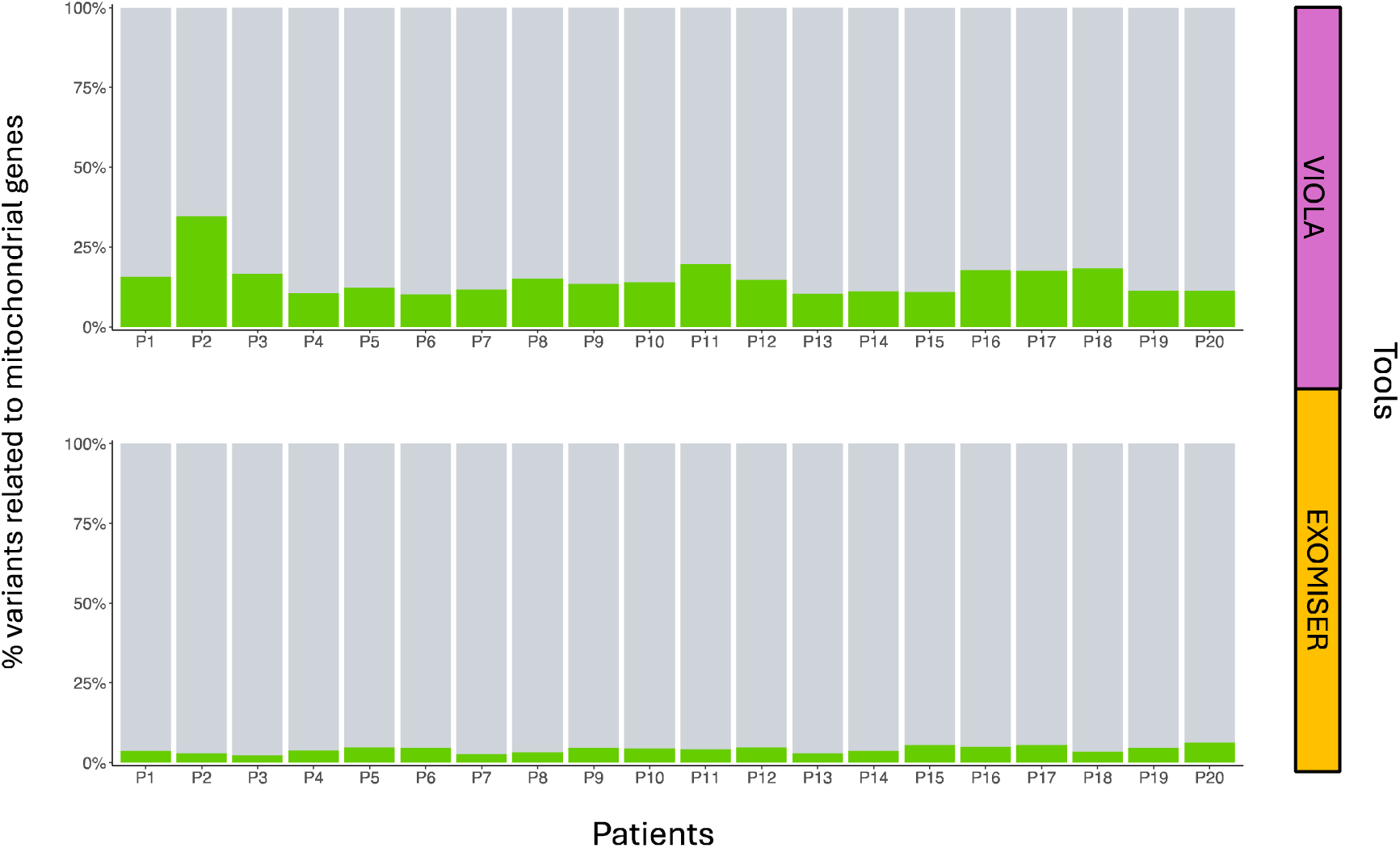
Proportion of candidate variants in mitochondrial genes prioritized by VIOLA and Exomiser. Stacked barplots show the percentage of candidate variants located in genes listed in the MitoCarta database for each of the 20 patients of the cohort. Each bar represents the distribution of variants prioritized by VIOLA (up) and Exomiser (down). Variants in mitochondrial genes are shown in green, while those in non-mitochondrial genes are in grey.

## Discussion and conclusion

In this study, we presented VIOLA, a novel tool for variant prioritization that relies on machine learning and multi-omics integration to help identify causal variants, in the particular context of MD. The combination of genomics, transcriptomics, and phenotype data in a comprehensive analytical framework allowed us to address some of the limitations inherent in existing variant prioritization methods.

One of VIOLA’s key steps is the combination of CADD scores via a VAE. The results of our analysis underline the importance of this step: shuffling CADD scores disrupted the accurate classification of causal variants in many cases, underscoring their essential role in the pipeline. We demonstrated the advantage to use VAE coupled with DBSCAN clustering compared to other technique, because it allowed us to achieve a robust identification of outlier variants. We showed the added value of incorporating transcriptomic data into the prioritization process. Through co-expression analysis, we identified gene modules enriched in mitochondrial genes, which significantly improved the relevance of variant ranking. The VIOLA ranking, comprising the Vscore and the VAscore proved effective in classifying causal variants. The integration of Exomiser and VIOLA scores into the VAscore demonstrated synergistic effects, consistently improving the classification of responsible variants across all patient cases tested. This reinforces the importance of combining complementary tools to achieve a more accurate prioritization of variants. Finally, we included an output easy to interpret in VIOLA pipeline. The special advantage of html format is the integration of direct links to databases widely used by physicians to assess the pathogenic potential of variants. By centralizing all relevant information in a single interactive file, we aim to simplify the process of identifying the causal variant responsible for a patient’s condition. This user-friendly result not only saves time, but also facilitates a more intuitive workflow, enabling geneticists to concentrate on interpreting results rather than navigating complex datasets.

Due to the heterogeneity of MDs, the difficulty of distinguishing between PDM and SMD, and the incomplete annotation of mitochondrial genes, some cases are particularly complex and have remained in diagnostic stalemate for decades. Some responsible variants may reside in genes whose link with mitochondria has not yet been established, and which are therefore not yet associated with MDs. Consequently, the causative variant(s) is (are) difficult to detect.

VIOLA successfully ranked the causal variants within the top five positions using the ARrank in all four diagnosed patients. Importantly, three of these variants were located in genes not currently annotated as mitochondrial, underlining VIOLA’s ability to identify novel disease mechanisms in the context of MD beyond known mitochondrial gene sets such as MitoCarta^36^ or the list of genes already known to be involved in MD^11^. This highlights the tool’s potential not only for accurate variant prioritization but also as an exploratory resource to uncover previously unknown factors in MD. VIOLA can therefore help diagnose complex or atypical cases, where conventional approaches may prove insufficient.

However, several challenges remain. The pathogenic variant can lie outside the coding regions of the genome. Since our data is based on WES, non-coding variants, which could have an impact on gene regulation or splicing, are not taken into account. In such cases, whole genome sequencing (WGS) would be necessary to obtain a more complete view of the patient’s variants, including those in non-coding regulatory regions^62^. However, the switch from WES to WGS poses additional problems. The large volume of WGS data considerably increases the complexity of the analysis, making it difficult to identify the responsible variant among thousands, if not millions, of potential candidates.

Future work should focus on extending the scope of the analysis to WGS data and incorporating functional validation experiments to confirm the pathogenicity of the highest-ranked variants. In addition, the weighting scheme for different Vscore features could be further optimized, possibly using larger and more diverse datasets.

In conclusion, VIOLA represents an exploratory tool to help diagnose complex cases of patients with suspected MD. By combining advanced computational methods with prior knowledge and different omics layer, VIOLA improves the identification of causal variants, paving the way for more effective diagnostics and personalized treatments for patients with suspected MD and potentially other rare diseases. Further development and validation of VIOLA on larger cohorts and diversified phenotypic datasets will be essential to establish its generalizability and robustness in different clinical and research settings.

---

**Supplementary Table I:** Number of retained variants at each step of the VIOLA pipeline for all patients. The number of remaining variants at each step of the VIOLA pipeline is reported in the table from the input file to the ranking with Autosomal Recessive rank (ARrank) for the 20 patients of the cohort. Abbreviations for steps are DBSCAN: Density-Based Spatial Clustering of Applications with Noise, HPO: Human Phenotype Ontology, Vrank: VIOLA rank, ARrank: Autosomal Recessive rank.

| Patient | Input | DBSCAN | Filters | HPO | Vrank | ARrank |
| --- | --- | --- | --- | --- | --- | --- |
| 1 | 77580 | 43103 | 18866 | 564 | 64 | 39 |
| 2 | 102136 | 55647 | 22935 | 649 | 78 | 59 |
| 3 | 90958 | 50128 | 20016 | 3559 | 375 | 272 |
| 4 | 171319 | 93011 | 42590 | 8603 | 882 | 739 |
| 5 | 152757 | 83319 | 37601 | 4252 | 455 | 378 |
| 6 | 15217 | 8025 | 3172 | 261 | 39 | 12 |
| 7 | 15946 | 8457 | 3397 | 370 | 51 | 18 |
| 8 | 15425 | 8144 | 3161 | 417 | 60 | 20 |
| 9 | 11633 | 5959 | 2387 | 349 | 52 | 20 |
| 10 | 51909 | 29157 | 12394 | 2444 | 266 | 174 |
| 11 | 78900 | 43915 | 19149 | 2695 | 295 | 228 |
| 12 | 70827 | 39695 | 18681 | 3488 | 388 | 274 |
| 13 | 80202 | 44685 | 18724 | 3890 | 428 | 299 |
| 14 | 85213 | 47266 | 20283 | 5406 | 580 | 434 |
| 15 | 82385 | 45851 | 19736 | 3677 | 387 | 275 |
| 16 | 85214 | 47248 | 20982 | 1064 | 124 | 87 |
| 17 | 84419 | 46767 | 20521 | 4070 | 441 | 303 |
| 18 | 51074 | 28675 | 12603 | 2229 | 252 | 188 |
| 19 | 80403 | 44825 | 19131 | 3268 | 357 | 259 |
| 20 | 97046 | 53121 | 23850 | 3373 | 407 | 340 |

## References

1. Nguengang Wakap, S., et al. Estimating cumulative point prevalence of rare diseases: analysis of the Orphanet database. Eur. J. Hum. Genet. 28, 165–173 (2020).

2. Haendel, M. et al. How many rare diseases are there? Nat. Rev. Drug Discov. 19, 77–78 (2020).

3. Marwaha, S., Knowles, J. W. & Ashley, E. A. A guide for the diagnosis of rare and undiagnosed disease: beyond the exome. Genome Med. 14, 23 (2022).

4. Gorman, G. S. et al. Mitochondrial diseases. Nat. Rev. Dis. Primer 2, 1–22 (2016).

5. Munnich, A. & Rustin, P. Clinical spectrum and diagnosis of mitochondrial disorders. Am. J. Med. Genet. 106, 4–17 (2001).

6. Niyazov, D. M., Kahler, S. G. & Frye, R. E. Primary mitochondrial disease and secondary mitochondrial dysfunction: importance of distinction for diagnosis and treatment. Mol. Syndromol. 7, 122–137 (2016).

7. Parikh, S. et al. Diagnosis of ‘possible’mitochondrial disease: an existential crisis. J. Med. Genet. 56, 123–130 (2019).

8. Mancuso, M. et al. International Workshop:: Outcome measures and clinical trial readiness in primary mitochondrial myopathies in children and adults. Consensus recommendations. 16–18 November 2016, Rome, Italy. Neuromuscul. Disord. 27, 1126–1137 (2017).

9. Hutchison III, C. A., Newbold, J. E., Potter, S. S. & Edgell, M. H. Maternal inheritance of mammalian mitochondrial DNA. Nature 251, 536–538 (1974).

10. Alston, C. L., Rocha, M. C., Lax, N. Z., Turnbull, D. M. & Taylor, R. W. The genetics and pathology of mitochondrial disease. J. Pathol. 241, 236–250 (2017).

11. Stenton, S. L. & Prokisch, H. Genetics of mitochondrial diseases: Identifying mutations to help diagnosis. EBioMedicine 56, (2020).

12. Wortmann, S. B., Koolen, D. A., Smeitink, J. A., van den Heuvel, L. & Rodenburg, R. J. Whole exome sequencing of suspected mitochondrial patients in clinical practice. J. Inherit. Metab. Dis. 38, 437–443 (2015).

13. Souche, E. et al. Recommendations for whole genome sequencing in diagnostics for rare diseases. Eur. J. Hum. Genet. 30, 1017–1021 (2022).

14. Dewey, F. E. et al. Clinical interpretation and implications of whole-genome sequencing. Jama 311, 1035–1045 (2014).

15. Ng, P. C. & Henikoff, S. Predicting deleterious amino acid substitutions. Genome Res. 11, 863–874 (2001).

16. Adzhubei, I. A. et al. A method and server for predicting damaging missense mutations. Nat. Methods 7, 248–249 (2010).

17. Kircher, M. et al. A general framework for estimating the relative pathogenicity of human genetic variants. Nat. Genet. 46, 310–315 (2014).

18. Smedley, D. et al. Next-generation diagnostics and disease-gene discovery with the Exomiser. Nat. Protoc. 10, 2004–2015 (2015).

19. Ioannidis, N. M. et al. REVEL: an ensemble method for predicting the pathogenicity of rare missense variants. Am. J. Hum. Genet. 99, 877–885 (2016).

20. Birgmeier, J. et al. AMELIE speeds Mendelian diagnosis by matching patient phenotype and genotype to primary literature. Sci. Transl. Med. 12, eaau9113 (2020).

21. Schubach, M., Maass, T., Nazaretyan, L., Röner, S. & Kircher, M. CADD v1.7: using protein language models, regulatory CNNs and other nucleotide-level scores to improve genome-wide variant predictions. Nucleic Acids Res. 52, D1143–D1154 (2024).

22. Hamilton, E. M. et al. UFM1 founder mutation in the Roma population causes recessive variant of H-ABC. Neurology 89, 1821–1828 (2017).

23. Chen, D.-H. et al. ADCY5-related dyskinesia: broader spectrum and genotype–phenotype correlations. Neurology 85, 2026–2035 (2015).

24. Rouzier, C. et al. Primary mitochondrial disorders and mimics: Insights from a large French cohort. Ann. Clin. Transl. Neurol. 11, 1478–1491 (2024).

25. Andrews, S. FastQC: a quality control tool for high throughput sequence data. *No Title* (2010).

26. McKenna, A. et al. The Genome Analysis Toolkit: a MapReduce framework for analyzing next-generation DNA sequencing data. Genome Res. 20, 1297–1303 (2010).

27. McLaren, W. et al. The Ensembl Variant Effect Predictor. Genome Biol. 17, 122 (2016).

28. Landrum, M. J. et al. ClinVar: public archive of relationships among sequence variation and human phenotype. Nucleic Acids Res. 42, D980–D985 (2014).

29. Landrum, M. J. et al. ClinVar: updates to support classifications of both germline and somatic variants. Nucleic Acids Res. 53, D1313–D1321 (2025).

30. Sherry, S. T., Ward, M. & Sirotkin, K. dbSNP—database for single nucleotide polymorphisms and other classes of minor genetic variation. Genome Res. 9, 677–679 (1999).

31. Chen, S. et al. A genomic mutational constraint map using variation in 76,156 human genomes. Nature 625, 92–100 (2024).

32. 1000 Genomes Project Consortium. A global reference for human genetic variation. Nature 526, 68 (2015).

33. Dobin, A. et al. STAR: ultrafast universal RNA-seq aligner. Bioinformatics 29, 15–21 (2013).

34. DeLuca, D. S. et al. RNA-SeQC: RNA-seq metrics for quality control and process optimization. Bioinformatics 28, 1530–1532 (2012).

35. Langfelder, P. & Horvath, S. WGCNA: an R package for weighted correlation network analysis. BMC Bioinformatics 9, 1–13 (2008).

36. Rath, S. et al. MitoCarta3. 0: an updated mitochondrial proteome now with sub-organelle localization and pathway annotations. Nucleic Acids Res. 49, D1541–D1547 (2021).

37. Kuleshov, M. V. et al. Enrichr: a comprehensive gene set enrichment analysis web server 2016 update. Nucleic Acids Res. 44, W90–W97 (2016).

38. Ashburner, M. et al. Gene Ontology: tool for the unification of biology. Nat. Genet. 25, 25–29 (2000).

39. The Gene Ontology Consortium et al. The Gene Ontology knowledgebase in 2023. Genetics 224, iyad031 (2023).

40. Gargano, M. A. et al. The Human Phenotype Ontology in 2024: phenotypes around the world. Nucleic Acids Res. 52, D1333–D1346 (2024).

41. Lachmann, A. et al. Geneshot: search engine for ranking genes from arbitrary text queries. Nucleic Acids Res. 47, W571–W577 (2019).

42. Hamosh, A., Scott, A. F., Amberger, J. S., Bocchini, C. A. & McKusick, V. A. Online Mendelian Inheritance in Man (OMIM), a knowledgebase of human genes and genetic disorders. Nucleic Acids Res. 33, D514–D517 (2005).

43. Grissa, D., Junge, A., Oprea, T. I. & Jensen, L. J. Diseases 2.0: a weekly updated database of disease–gene associations from text mining and data integration. Database 2022, baac019 (2022).

44. Landrum, M. J. et al. ClinVar: improving access to variant interpretations and supporting evidence. Nucleic Acids Res. 46, D1062–D1067 (2018).

45. Siepel, A. et al. Evolutionarily conserved elements in vertebrate, insect, worm, and yeast genomes. Genome Res. 15, 1034–1050 (2005).

46. Abascal, F. et al. Expanded encyclopaedias of DNA elements in the human and mouse genomes. Nature 583, 699–710 (2020).

47. Cheng, J. et al. MMSplice: modular modeling improves the predictions of genetic variant effects on splicing. Genome Biol. 20, 48 (2019).

48. Taliun, D. et al. Sequencing of 53,831 diverse genomes from the NHLBI TOPMed Program. Nature 590, 290–299 (2021).

49. Abadi, M., et al. TensorFlow: Large-scale machine learning on heterogeneous systems. (2015).

50. Sander, J., Ester, M., Kriegel, H.-P. & Xu, X. Density-based clustering in spatial databases: The algorithm gdbscan and its applications. Data Min. Knowl. Discov. 2, 169–194 (1998).

51. Rahmah, N. & Sitanggang, I. S. Determination of optimal epsilon (eps) value on dbscan algorithm to clustering data on peatland hotspots in sumatra. in vol. 31 012012 (IoP Publishing, 2016).

52. De Maesschalck, R., Jouan-Rimbaud, D. & Massart, D. L. The Mahalanobis distance. Chemom. Intell. Lab. Syst. 50, 1–18 (2000).

53. R Core Team, R. R: A language and environment for statistical computing. (2013).

54. Stelzer, G. et al. The GeneCards suite: from gene data mining to disease genome sequence analyses. Curr. Protoc. Bioinforma. 54, 1–30 (2016).

55. GTEx Consortium et al. The Genotype-Tissue Expression (GTEx) pilot analysis: multitissue gene regulation in humans. Science 348, 648–660 (2015).

56. Dyer, S. C. et al. Ensembl 2025. Nucleic Acids Res. 53, D948–D957 (2025).

57. Wold, S., Esbensen, K. & Geladi, P. Principal component analysis. Chemom. Intell. Lab. Syst. 2, 37–52 (1987).

58. Pedregosa, F. et al. Scikit-learn: Machine learning in Python. J. Mach. Learn. Res. 12, 2825– 2830 (2011).

59. Van der Maaten, L. & Hinton, G. Visualizing data using t-SNE. J. Mach. Learn. Res. 9, (2008).

60. Kremer, L. S. et al. Genetic diagnosis of Mendelian disorders via RNA sequencing. Nat. Commun. 8, 15824 (2017).

61. Murdock, D. R. et al. Transcriptome-directed analysis for Mendelian disease diagnosis overcomes limitations of conventional genomic testing. J. Clin. Invest. 131, (2021).

62. Mattick, J. S., Dinger, M., Schonrock, N. & Cowley, M. Whole genome sequencing provides better diagnostic yield and future value than whole exome sequencing: The integration of genome sequencing with clinical records and data from the internet of things will transform health care. Med. J. Aust. 209, (2018).

